# Optimised CRISPR-Cas9 mediated single molecule imaging for accurate quantification through endogenous expression

**DOI:** 10.1101/482596

**Authors:** Abdullah O. Khan, Carl W. White, Jeremy A. Pike, Jack Yule, Alexandre Slater, Stephen J. Hill, Natalie S. Poulter, Steven G. Thomas, Neil V. Morgan

**Affiliations:** Institute of Cardiovascular Science, College of Medical and Dental Sciences, University of Birmingham, Edgbaston, B15 2TT; Centre of Membrane and Protein and Receptors (COMPARE), University of Birmingham and University of Nottingham, Midlands, UK; Division of Physiology, Pharmacology and Neuroscience, School of Life Sciences, University of Nottingham, Nottingham; Molecular Endocrinology and Pharmacology, Harry Perkins Institute of Medical Research, Nedlands, WA, Australia; Centre for Medical Research, The University of Western Australia, Crawley, WA, Australia

**Keywords:** Single molecule imaging, super-resolution microscopy, CRISPR-Cas9 genome editing, CXCR4, receptor biology, cytoskeleton

## Abstract

The use of CRISPR-Cas9 genome editing to introduce endogenously expressed tags has the potential to address a number of the classical limitations of single molecule localisation microscopy. In this work we present the first systematic comparison of inserts introduced through CRISPR-knock in, with the aim of optimising this approach for single molecule imaging. We show that more highly monomeric and codon optimised variants of mEos result in improved expression at the *TubA1B* locus, despite the use of identical guides, homology templates, and selection strategies. We apply this approach to target the G protein-coupled receptor (GPCR) CXCR4 and show a further insert dependent effect on expression and protein function. Finally, we show that compared to over-expressed CXCR4, endogenously labelled samples allow for accurate single molecule quantification on ligand treatment. This suggests that despite the complications evident in CRISPR mediated labelling, the development of CRISPR-PALM has substantial quantitative benefits.

---

Super-resolution imaging techniques offer unique insights into cellular structures and organisations on a nanoscale inaccessible through conventional light microscopy. Of the many super-resolution methods currently available, single molecule localisation microscopy (SMLM) is unique in that it not only offers spatial resolution in the tens of nanometers(1–6), but a means of quantifying single emitters for insights into protein stoichiometry and organisation(2–4, 7).

PALM (Photo Activated Localisation Microscopy) and STORM (Stochastic Optical Reconstruction Microscopy) are archetypal SMLM methods which are based on the stochastic activation of subsets of molecules within a sample to allow for precise localisation and rendering post-acquisition(2, 3). The key difference between these two approaches lies in the nature of the reporters used to generate ‘blinking’ events which result from the stochastic activation of fluorophores. While in STORM paired dyes (nSTORM) or organic fluorophores capable of switching within an appropriate redox buffer (dSTORM) are used, PALM relies on photoswitching fluorescent proteins (e.g.mEos,Dendra). Since the early inception of these techniques, a number of variations have been developed which aim to address the shortcomings of previous iterations and improve on both spatial resolution and the quantitative capability of SMLM (including iPALM, PAINT, and MINIFLUX)(8–10).

SMLM techniques are potentially potent quantitative tools, however single molecule imaging and quantification have important caveats which are critical for accurate image acquisition, reconstruction, and quantification. First and foremost is the correct identification of single molecules, as the phenomenon of ‘blinking’ involves repeated cycling between on-off states, and persistence across multiple frames such that ‘single’ molecules are more often collections of peaks (11–13). This is a significant limitation in single molecule quantification, and a considerable focus of the field is accurately detecting individual emitters and modelling fluorophore behaviour.

A significant advance in the field would be the development of systems with a known labelling stoichiometry and known photoswitching kinetics. While PALM addresses both to an extent by introducing photoswitchable tags with a known 1:1 labelling ratio, the variability and artefacts introduced by over-expression are one of a number of problems inherent to this approach(14). The endogenous expression of labels can offer a solution to this particular quantitative caveat of SMLM, and offers a number of other distinctive benefits. Cells expressing a genomically encoded tag will provide a large sample with defined expression profiles and label distributions, thus minimising sample to sample variation which is a major concern in nanoscale measurements(15, 16).

This approach has been used very successfully in prokaryotic cells, with a number of excellent studies highlighting the utility of endogenous labels in applying SMLM to interrogate nanoscale cellular architecture(17–19). The application of this approach in mammalian cells has been limited in the past due to the poor efficiency and high costs associated with mammalian gene editing. With the advent of CRISPR-Cas9 gene editing however, the insertion of labels into target endogenous loci has become substan-tially more convenient(20–22). To date a handful of studies have applied this approach in live and fixed cell super resolution microscopy, and of these only two studies use the integration of photoswitchable tags for CRISPR mediated SMLM(15, 16, 23, 24). Cho *et al.* do not report on the effect of knock-in on expression level, or the resulting quality of their single molecule imaging. Hansen *et al.* effectively used their HaloTag knock-ins, and while the data is not shown, report the generation of homozygous knockins with no effect of insertion on the target’s expression level compared to wild type protein(23, 24).

In our previous work, we report a tag dependent expression of CRISPR edited *TubA1B* in multiple cell lines. While mEGFP is sufficiently expressed in Hek293T, A549, and Hel 92.1.7 to allow for diffraction limited imaging, mEos 3.2 tagged cells in all three samples show significantly reduced expression when compared to mEGFP knock-in. Of the three cell lines tested, only Hel 92.1.7 cells express mEos 3.2 tagged *TubA1B* at a high enough level to generate images comparable to dSTORM, according to quantitative measurements of resolution. As only heterozygous knock-ins were found, we reasoned that the expression of tagged tubulin was regulated to maintain cell function and morphology, and that where the intrinsic qualities of fluorophores have a more dramatic effect on protein function, the tagged allele would be subsequently down-regulated. As mEGFP is a highly monomeric, codon optimised fluorophore, its expression is better tolerated than mEos 3.2. In this work we evaluate the effect of optimising a fluorescent protein on the expression of CRISPR-Cas9 generated knock-in cells (Figure 1). We test mEos variants with improved monomericity and codon optimisation, and show a substantial improvement on knock-in to the *TubA1B* locus. To determine whether this approach is suitable for receptor quantification, we apply this approach to target CXCR4, a GPCR with significant therapeutic value, to evaluate the effect of knock-ins on receptor distribution and function. Finally, we explore the quantitative potential of CRISPR knock-in when compared to over-expression. We report a tag and gene dependent effect of insertion with significant implications for both single molecule imaging and the wider field of CRISPR mediated HDR (homology directed repair). To our knowledge, the behaviours of different tags in identical knock-ins have not been reported or systematically studied. Similarly we show that compared to over-expression of CXCR4, CRISPR knock-in labelling of this receptor allows for accurate single molecule clustering analysis. This is a significant advance in the field of imaging receptor behaviours on the nanoscale, suggesting that despite the complications inherent to knock-in design, CRISPR-PALM is a powerful quantitative single molecule tool.

**Fig. 1.**
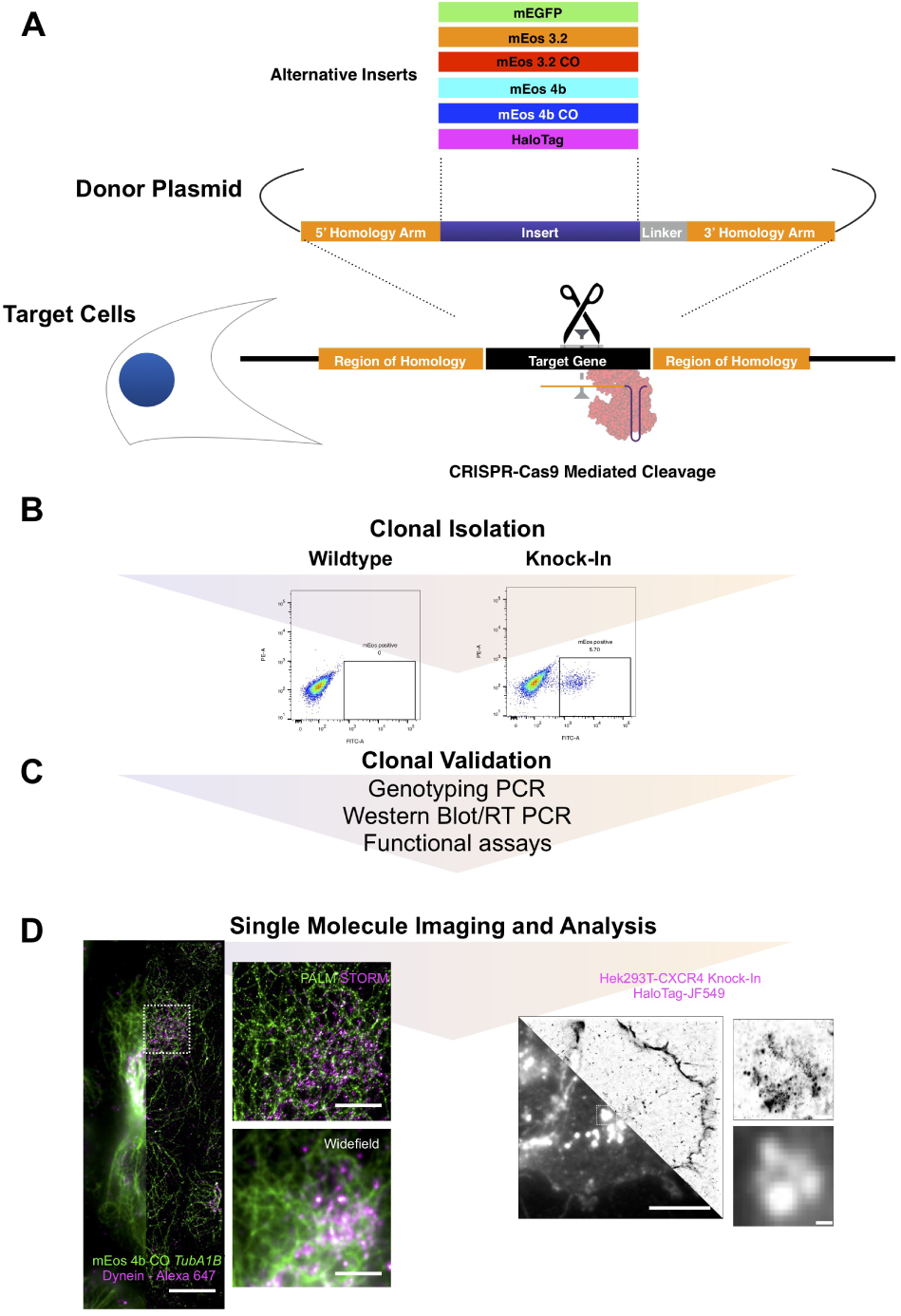
Summary of experimental workflow. (A) Multiple inserts including optimised mEos variants were cloned into identical donor plasmids to assess the effects of fluorophore properties on endogenous expression after CRISPR knock-in. (B) Cells were identically transfected with specific guides, and after the clonal isolation of the brightest (and hence best expressing cells) by single cell sorting, (C) cells were validated for insertion efficiency, expression level, and functional effects. (D) Finally successful clones were interrogated for their efficiency as tools for single molecule microscopy through measurements of effective resolution (*TubA1B)* and cluster distribution (*CXCR4*).

## Results

### Monomeric and codon optimised variants increase endogenous expression at the *TubA1B* locus for improved single molecule imaging

We reasoned that the variation in CRISPR labelled *TubA1B* observed in previous work was due to fundamental differences in fluorophore properties which could lead to down-regulation or degradation. mEGFP is a highly monomeric, codon optimised fluorescent protein while mEos 3.2 has been reported to oligomerise(15, 25). To test this hypothesis, HDR donor templates for *TubA1B* were generated carrying a codon optimised mEos 3.2 and both the original and a codon optimised version of mEos 4b (Figure S1) reported by Paez-Segala *et al.* (26). These donors were co-transfected with a *TubA1B* targeting guide and Cas9 expressing plasmid into Hel 92.1.7, a cell line which we previously demonstrated expresses high levels of *TubA1B* but still exhibits significantly less mEos 3.2 tagged *TubA1B* when compared to endogenously labelled mEGFP at the same locus(15). As in our previous work, cells were single sorted to select for the most fluorescent clones (and therefore cells which best express a specific label) (Figure S2A). Individual clones were expanded and further validated by flow assisted cell sorting (FACS) and immunofluorescence to ensure adequate expression before valida-tion by PCR (Figure S2B). As previously reported by ourselves and Roberts *et al.*, no homozygous expression of tagged *TubA1B* is observed (loss of the genomic wild type band and wild type protein). Interestingly however, markedly different expression profiles are evidenced on western blotting of validated clones (Figure 2A). In wild type cells one band at the predicted molecular weight of tubulin is observed (approx. 50 kDa), while in each of our CRISPR knock-ins a heavier band consistent with the fusion protein is observed and quantified (approx. 80kDa) (Figure 2A). When compared to our previously reported mEos 3.2 clones, no significant difference in expression is observed in clones carrying a codon optimised variant of the original mEos 3.2 (Figure 2B). mEos 4b shows a significant decrease in expression (p< 0.0001), however the codon optimised version of this sequence demonstrates a significant increase in expression when compared to the mEos 3.2 clone (p = 0.0005, 0.0244, and 0.022 respectively) (Figure 2B). The highest expressing mEos 4b CO clone was taken forward to evaluate the effect of improved label expression on the resulting single molecule imaging. PALM imaging of clone C4 results in high quality single molecule images which demonstrate a significantly increased number of localisations compared to mEos 3.2 (p= 0.0230) (Figure 2D). No significant difference in other measures of resolution, namely Fourier Ring Correlation and localisation precision (XY uncertainty) were observed (Figure 2E,F).

**Fig. 2.**
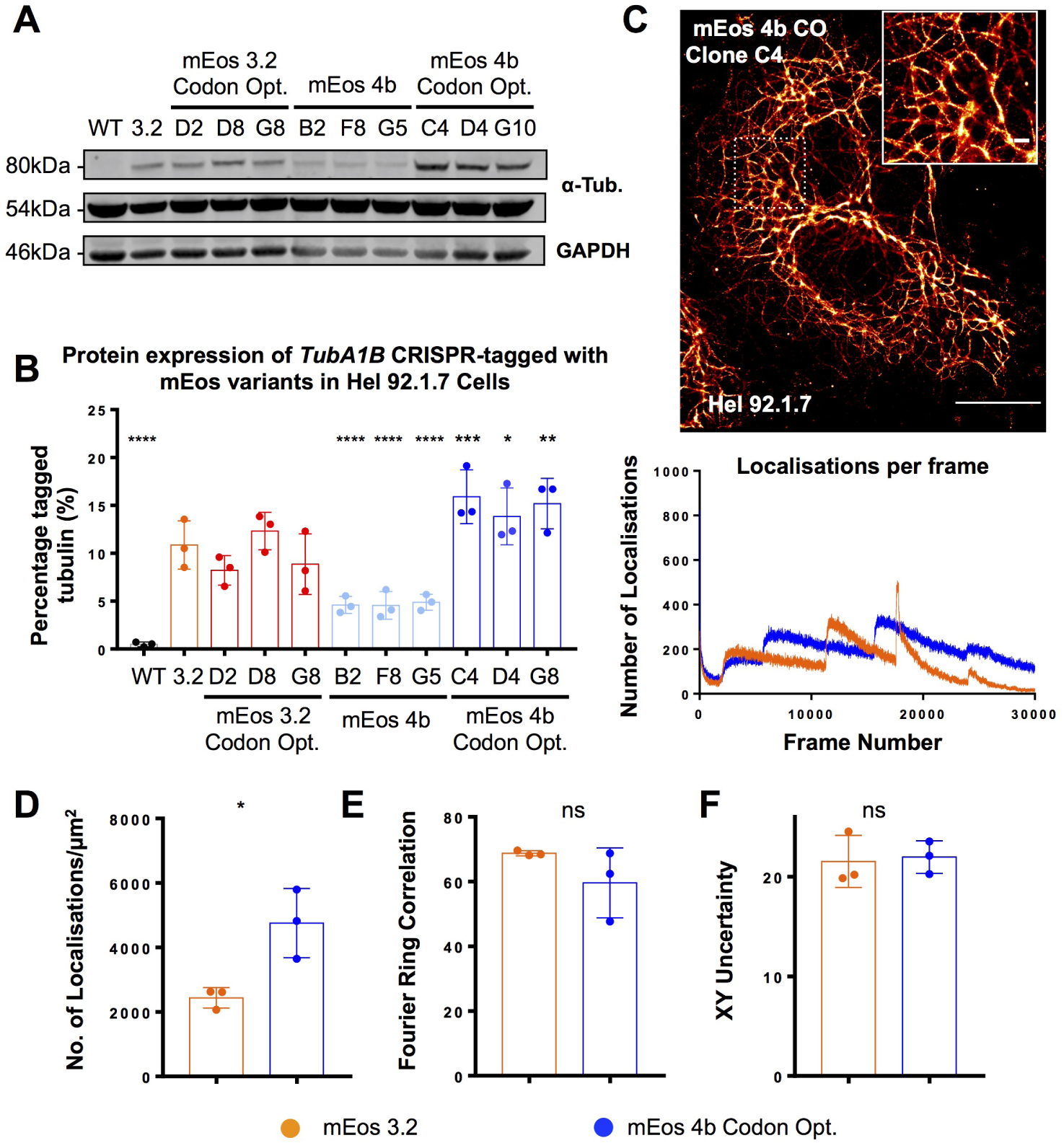
Optimised expression of mEos significantly improves SMLM imaging. Hel 92.1.7 clones CRISPR edited to express mEos 3.2, codon optimised mEos 3.2, mEos 4b, and codon optimised mEos 4b TubA1B inserts were isolated and compared to a previously reported *TubA1B*-mEos 3.2 clone to determine the effects of fluorophore properties and codon optimisation on expression level. (A) Western blotting of clones carrying specific mEos variants at the TubA1B locus showed a variation in expression of tagged tubulin dependent on the insert. (B) No significant difference was observed in mEos 3.2 codon optimised clones, however a significant reduction in expression was observed in cells carrying mEos 4b. Conversely, a significant improvement in expression was observed in clones carrying a codon optimised mEos 4b. (C,D) mEos 4b CO clones produce high quality PALM images, with a significantly increased number of localisations compared to mEos 3.2 clones. (E,F) No significant difference in fourier ring correlation or localisation precision were observed. *n* = 3 (S.D), statistical analysis performed using One-Way ANOVA with multiple comparisons to the mEos 3.2 clone for western data, and by t-test for quantitative SMLM comparisons.

In our previous work we reported a significant down-regulation of *TubA1B*-mEos 3.2 when compared to mEGFP in Hek293T cells, and show that in these knock-in cell lines the poor expression of mEos 3.2 labelled cells does not allow for the clear resolution of microtubules and performs poorly when quantitatively compared to dSTORM images of Hek293T(15). To determine whether the improvement in expression achieved by mEos 4b CO knock-in improves the quality of SMLM in Hek293T, where *TubA1B* is more poorly expressed compared to Hel 92.1.7, we generated Hek293T clones carrying *TubA1B*-mEos 4b CO and confirm the heterozygous insertion of the tag before quantitatively establishing expression (Figure S3). Compared to previously generated mEos 3.2 clones, two of the three mEos 4b CO tagged cells show a significant improvement in expression (Figure 3A). While this improvement is not as substantial as the increase observed in Hel 92.1.7 and remains significantly lower than mEGFP expression, there is a significant effect on the quality of resulting single molecule images (Figure 3B). This is quantified as a significant improvement in the number of localisations, FRC, and localisation precision (XY uncertainty) (Figures 3B), indicating that across both cell lines observed the codon optimised, monomeric mEos variant is better expressed, and the resulting improvement in the distribution of the endogenously expressed label also improves SMLM resolution and performance.

**Fig. 3.**
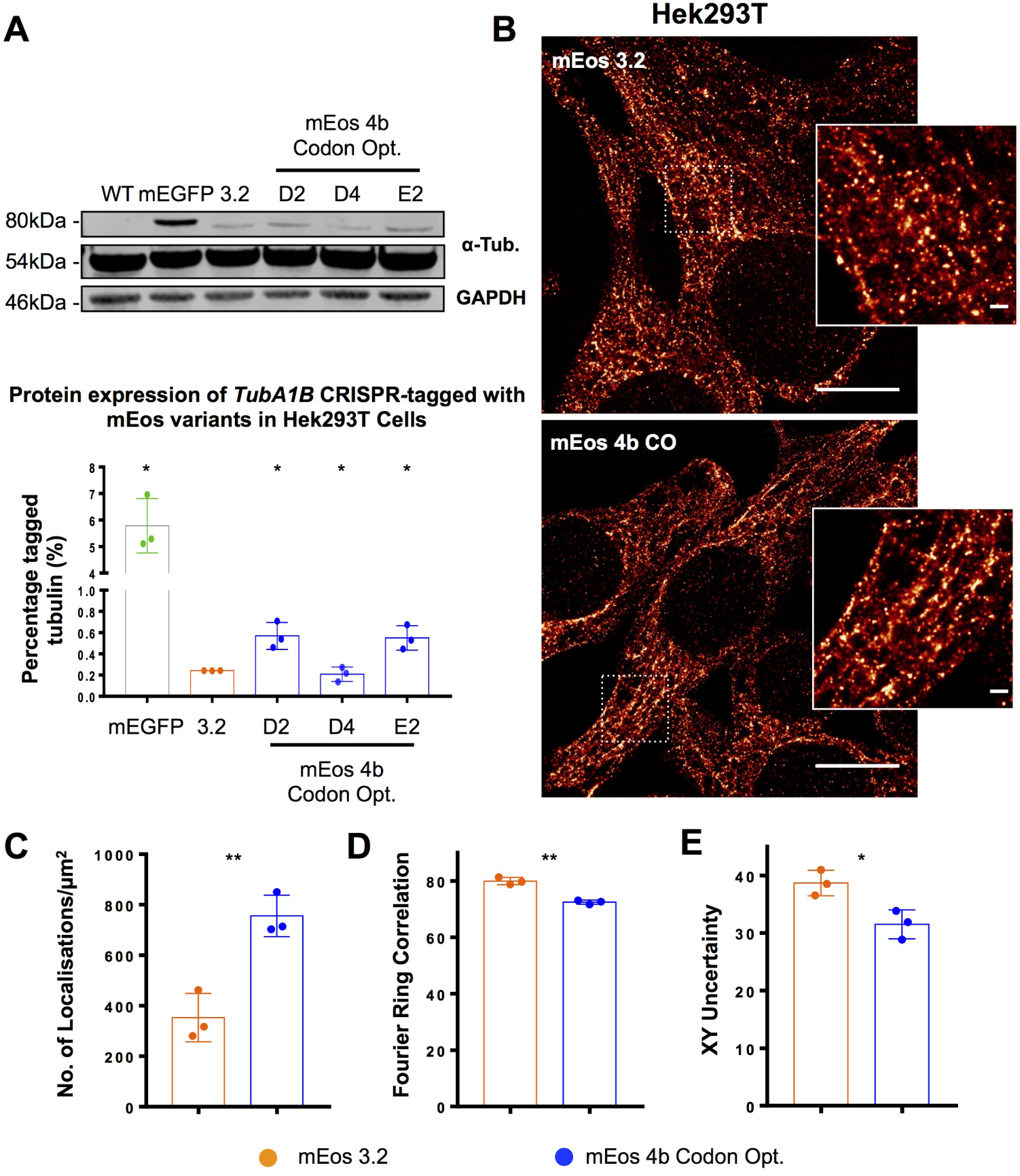
Knock-in of codon optimised mEos 4b in Hek293T results in a 2-fold improvement in expression compared to mEos 3.2. (A) mEos 4b CO knock-in Hek293T were generated and compared to previously reported mEGFP and mEos 3.2 knock-ins. A 2-fold, significant improvement in expression is observed in western blotting when compared to mEos 3.2 (* p = 0.0473, ns p = 0.49, * p = 0.0444 clones D2, D4, and E2 respectively). (B) mEos 4b CO labelled cells demonstrate improved microtubule labelling, which is quantifiable and observed as an improvement in (C) the number of localisations (** p = 0.0053), (D) FRC (** p = 0.0010), and (E) modal XY uncertainty (* p = 0.0210) when compared to mEos 3.2. (*n* = 3 (S.D), Western Blotting tested by 2-way ANOVA with multiple comparisons, SMLM quantifications by t-testing.)

### CRISPR targeting of CXCR4 allows for diffraction limited and super-resolved imaging of knock-in tagged receptor

Beyond its capacity for nanoscale resolution, single molecule imaging allows for unique quantification strategies which provide insights into the distribution, clustering behaviours, and stoichiometry of proteins of interest. Information of this kind is particularly relevant to the study of receptors, and as such the generation of a single molecule reporter based on endogenous expression is of particular interest to the field of receptor biology. The C-X-C chemokine receptor type 4 (CXCR4) CXCR4 is a G protein-coupled receptor essential for the proliferation of B-cell precursors during haematopoiesis, and plays a role in the homing and retention of most immune cells(27–29). It is also vital for the development of the hematopoietic, cardiovascular, and nervous systems during embryogenesis(30–34). Dysregulation of CXCR4 function can result in cancer progression and metastasis, immunodeficiency diseases (particularly WHIM syndrome which is characterised by mutations in CXCR4), neurodevelopmental defects and CXCR4 acts as a co-receptor in facilitating HIV infection. CXCR4 has therefore been the focus of major drug discovery efforts. CXCR4 forms homo- and heterooligomers as well as receptor clusters which affect its signalling properties, making it an ideal target for proof-of-principle CRISPR-PALM knock-in studies to establish receptor kinetics and stoichiometry in response to novel compounds(35–38). CXCR4 also presents a number of labelling challenges which are addressed using a gene-editing approach, namely poor labelling by antibodies for both immunofluorescence (and hence dSTORM) and western blotting (Figures 4A, S4) which results in a dependence on over-expression systems to study the receptor. CRISPR knock-in has successfully ameliorated the absolute need for over-expression of donor fused CXCR4 in the bioluminescence resonance energy transfer assay as demonstrated by White *et al.*, suggesting that C-terminal knock-ins are functional for this receptor. Knock-in templates, described previously by White *et al.*, were designed to target CXCR4 and introduce mEGFP, mEos 4b CO, and a codon optimised HaloTag sequences to the C-terminus. HaloTag was included to expand the repertoire of single molecule labels and due to the range of organic fluorophores available for the labelling of this tag, allows for the potential future multiplex with existing fluorescent proteins in CRISPR-PALM studies. As a chemical tag which is non-fluorescent until exposed to an appropriately modified organic fluorophore, HaloTag offers the benefits of a one to one labelling ratio through endogenous expression as well as the high photon counts (and thus localisation precision) offered by organic fluorophores. In cells expressing CXCR4-mEGFP or CXCR4-HaloTag knock-ins, labelled protein was observed as expected at both the plasma membrane and endosomal sites, surprisingly however, we found that mEos 4b CO appeared to form aggregates in PCR validated clones (Figures 4A, S5), either due to mis-localisation of the fusion protein or cleavage of the tag. mEGFP and HaloTag knock-ins demonstrate even labelling at the plasma membrane across the clonal population, (Figure 4B), suggesting that knock-in establishes a stably expressing population suitable for large sample sizes.

**Fig. 4.**
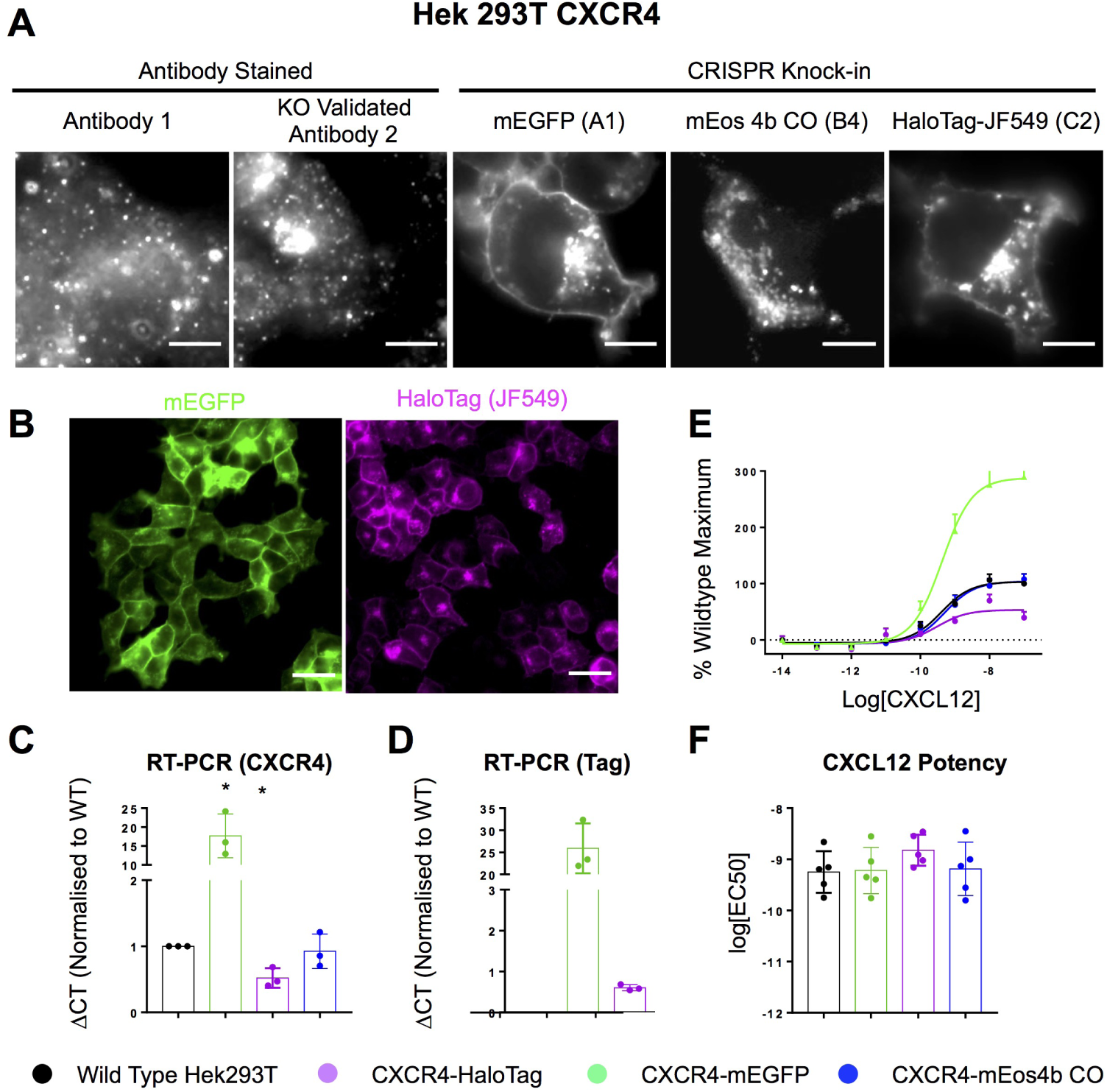
CRISPR-Cas9 mediated knock-in at the CXCR4 C-terminus results in endogenously labelled protein which localises to the membrane and endosomes. (A) Staining of CXCR4 with KO validated and Fusin antibodies results in poor labelling of the receptor protein, while mEGFP and HaloTag knock-in cells demonstrate a membrane specific localisation. Interestingly mEos 4b CO clones demonstrate fluorescent aggregates, suggesting the fluorophore is cleaved. (B) mEGFP and HaloTag knock-in samples provide clonal populations which are evenly labelled and offer large samples for imaging experiments. (C) Measurement of total CXCR4 expression in wild type versus CRISPR edited cells through qRT-PCR. Compared to wild type, mEGFP knock-ins demonstrate an approximately 20-fold increase in expression (* p = 0.0384). Conversely, HaloTag knock-ins demonstrate a 50% reduction in expression (* p = 0.0301), while mEos 4b CO knock-in shows no change. (D) When RT PCR is performed using tag specific primers, the significant increase in mEGFP expression is likely due to the stability of mEGFP tagged CXCR4 RNA, suggesting that RNA stability is likely a factor in mediating the tag specific down-regulation observed. (E) CXCL12-mediated g protein activation measured by observing G protein dissociation/conformational changes by BRET in knock-in or wildtype cells transfected with cDNA encoding G*α* i1/Nluc and Venus/G*γ* 2. Results are expressed as % maximal CXCL12 response observed in wildtype cells andshow a 3 fold increase in response in mEGFP knock-in clones compared to wild type, with a 50% reduction in response in HaloTag knock-in, and no variation in mEos 4b CO clones. (F) Potency of CXCL12-mediated g protein activation in cells expressing wildtype or gene-edited CXCR4.(*n = 3* (S.D) for qRT-PCR data, compared by One-Way ANOVA with multiple comparisons. *n = 5* for functional data.)

Due to poor specific detection by western blotting, expression was determined by quantitative real-time PCR (qRT-PCR) (Figure S5). Compared to wild type, CXCR4- mEGFP knock-in clones demonstrate a substantial increase in expression (approximately 20 fold increase, (* p = 0.038)), while CXCR4-HaloTag knock-ins have an approximately 50% reduction in expression (Figure 5C). No significant change in CXCR4 expression was observed in cells expressing the CXCR4-mEos 4b CO knock-ins, which when paired with CXCL12 induced signalling responses that are comparable to wildtype suggests that the tag is cleaved with no impairment on function or expression (Figure 5C, E, F). RT-PCR using primers specific to tagged CXCR4 demonstrate that the increased expression of CXCR4-mEGFP is likely due to an increase in mRNA stability, suggesting that mRNA stability is likely a factor influencing the variations in expression observed. Furthermore, as reported by White *et al.*, the addition of tags to the C-terminus of CXCR4 was well tolerated with no differences in the potency of CXCL12-mediated G-protein activation (Figure 4F) or modulation of cAMP production (FigureS6) between cells expressing wild type CXCR4 or gene-edited tagged CXCR4. However in agreement with an increase in CXCR4-mEGFP RNA expression, an increase in the maximal response elicited by CXCL12 was observed (Figure 4E) indicating an increase in tagged receptor expression.

**Fig. 5.**
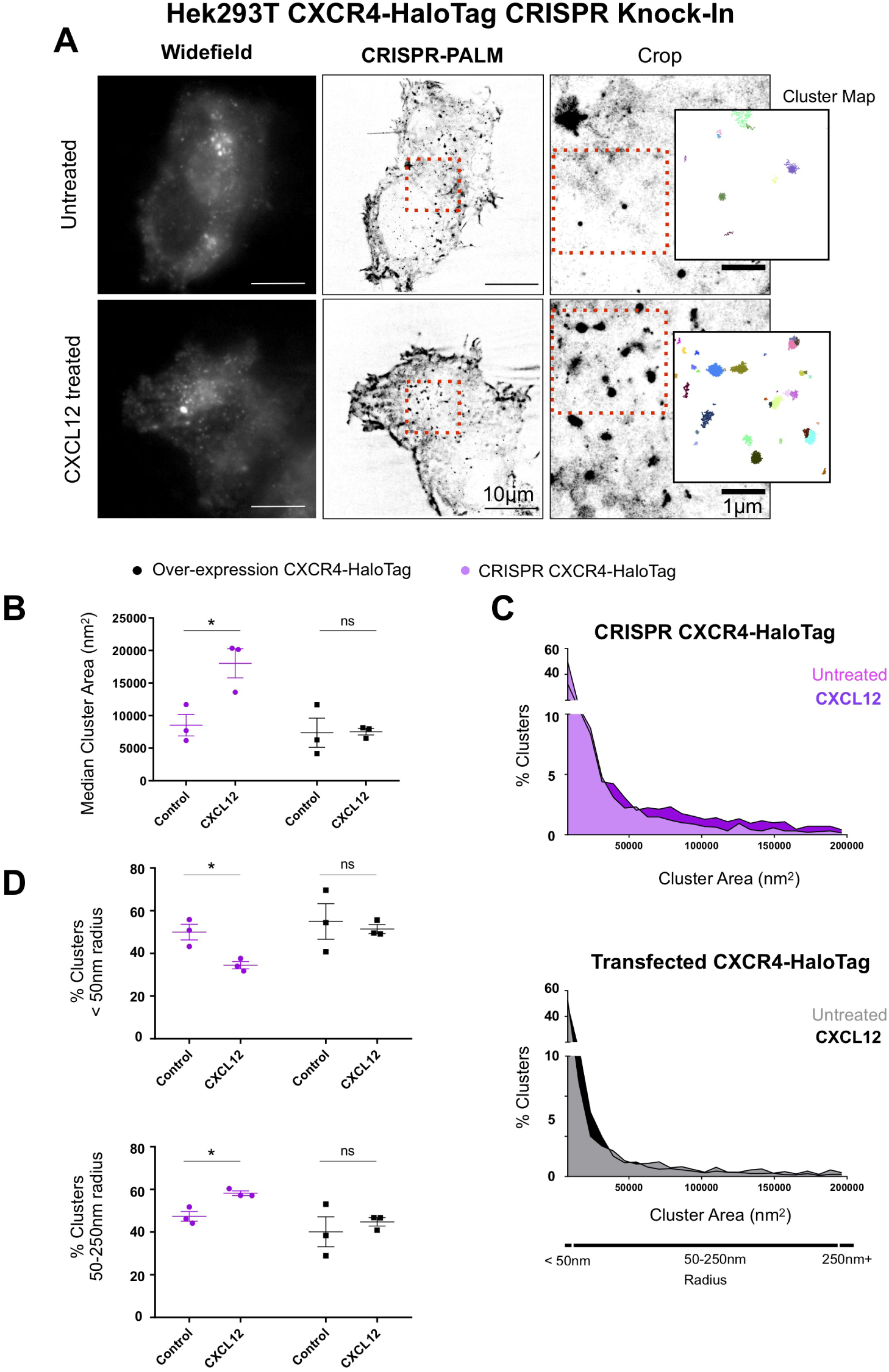
CRISPR Knock-In CXCR4 clones allow for accurate single molecule cluster analysis compared to transfected cells. (A) Single molecule imaging of HaloTag CXCR4 Knock-Ins allows for effective single molecule localisation microscopy through labelling with Janelia Fluor 549, and subsequent cluster analysis using persistence based clustering. (B) CXCR4 knock-in and Hek293T transfected with a CXCR4-HaloTag were subject to single molecule imaging and cluster analysis after treatment with CXCL12 (10*μ*M). CRISPR knock-ins demonstrate a significant increase in median cluster area on ligand treatment (* p = 0.0272), while transfected cells show no significant change.(C) Histograms showing percentage of clusters across a range of areas show that while a substantial shift towards larger cluster sizes is evident in CRISPR knock-ins, this is reduced in transfected cells. (D) The percentage of clusters below 50nm in radius is significantly reduced in CRISPR cells (* p = 0.0384) upon treatment with CXCL12 (upper panel), with a parallel increase in the percentage of clusters of larger sizes consistent with endocytic trafficking observed (* p = 0.0133) (lower panel). Conversely, no significant change in either collection of cluster sizes is evident in transfected cells. (*n* = 3, S.E.M). paired t-test.)

Finally, the most significant implication of developing knock-in lines for single molecule imaging lies in the generation of a robust, consistently labelled sample which allows for the quantitative analysis of receptor organisation and function. To interrogate whether CRISPR generated CXCR4-HaloTag knock-ins offer a means of robust single molecule quantitation, knock-in cell lines were compared to Hek293T transfected with a CXCR4-HaloTag over-expression vector after treatment with CXCL12. Labelling CXCR4-HaloTag with Janelia Fluor 549 (JF549) allows for stochastic activation and subsequent single molecule detection and cluster analysis (Figure 5A).

Compared to Hek293T over-expressing CXCR4-HaloTag, CRISPR Knock-In cells demonstrate a significant increase in median cluster area upon ligand treatment (* p = 0.0272) (Figure 5B). Conversely, transfected cells do not show a statistically significant difference in size (p = 0.9484). Histograms plotting the percentage of clusters by area shows a clear shift towards larger clusters on ligand treatment in CRISPR knock-in cells, but no clear trend in transfected cells. This suggests that heterogeneity in over-expressing samples is likely responsible for the lack of a significant effect on ligand treatment(Figure 5C).

A significant reduction in the percentage of clusters with a radius below 50nm is observed in CRISPR cells (* p = 0.0384), alongside a parallel increase in the percentage of clusters with radii between 50-250nm (* p = 0.0133). No such trend is observed in transfected samples (p = 0.7277, p = 0.57557 respectively). These results are consistent with the trafficking and clustering of CXCR4 on ligand treatment, and show that while variations in transfected samples limit the application of quantitative single molecule imaging, CRISPR knock-in cells offer a consistently labelled sample which allows for the quantitative measurement of changes in cluster size and distribution on ligand treatment.

## Discussion

In this study, we expand on our previous observation of a tag dependent effect on the expression of CRISPR labelled genes. In our original work, we hypothesised that the reduction in expression observed in mEos 3.2 tagged *TubA1B* when compared to equivalent mEGFP knockins is likely due to fundamental properties of the fluorophore affecting function, and therefore driving a regulatory down-regulation of the targeted allele. We expand on the original observation by testing mEos variants with improved monomericity and codon optimisation to determine whether, where identical selection strategies, CRISPR guides and donor templates are used, variations in expression attributable to these properties are observed. In Hel 92.1.7 cells, which highly express the *TubA1B* isoform and demonstrate sufficient labelling through mEos 3.2 expression for high quality CRISPR PALM, the introduction of a codon optimised, monomeric variant (mEos 4b CO) results in a significant improvement in expression as shown by western blotting, and as a result, an improvement in the number of single molecule detections recorded in subsequent imaging experiments. Codon optimisation of the original mEos 3.2 sequence alone was not enough to improve expression, similarly the original mEos 4b sequence was more poorly expressed. These findings led us to conclude that improved monomericity and codon optimisation together improve expression in Hel 92.1.7, and counter-acts the down regulation of mEos labelled *TubA1B* we had previously observed. To expand on this finding we generated equivalent CRISPR knock-ins in Hek293T, a cell line which demonstrates a significant down-regulation of the mEos tagged *TubA1b* gene in our previous work.

The original mEos 3.2 *TubA1B* clones result in extremely poor SMLM imaging. mEos 4b CO Hek293T knock-ins demonstrate a 2-fold increase in expression which results in a substantial improvement in quantitative SMLM metrics, however expression of the mEos 4b CO tagged *TubA1B* remains significantly less than mEGFP labelling of the same locus. This finding suggests that there are likely other properties of both fluorophore and sequence impeding expression in Hek293T, which may include maturation rates or further regulation at the RNA level. To investigate the applicability of this CRISPR-mediated labelling approach to the study and quantification of membrane receptors, we generated Hek293T cells carrying mEos 4b CO, mEGFP, and HaloTag knock-ins at the C-terminus of CXCR4 following the design of White *et al.* in their work establishing a Nanoluc knock-in at the same locus. CXCR4 is an ideal target for CRISPR approaches due to poor antibody labelling and the importance of receptor dynamics in viable drug targets like GPCRs. We included a codon optimised HaloTag sequence as self-labelling enzymes are extensively used in over-expression studies of GPCRs, and these approaches offer substantial imaging benefits (including the possibility of multiplexing with mEos, high photon counts and a plethora of conjugated labels). Interestingly we found a series of different expression and functional profiles on knock-in. mEos 4b CO appears to be cleaved and aggregates within the cell, with no evidence of functional effects on the receptor. mEGFP knock-ins label the receptor well, however a functional over-expression is observed which is likely due to the elevated stability of mEGFP-CXCR4 RNA as evidenced by qRT-PCR. Finally, HaloTag knock-ins result in the only homozygous insertion observed in our studies thus far, suggesting that this tag is well tolerated. However a 50% reduction in expression is observed in these knock-ins, as well as a similar reduction in CXCL12 response. Recently Hansen *et al.* report the use of HaloTag knock-in lines in the study of chromatin loop stability(24). The authors report the generation of homozygous knock-ins which express HaloTag labelled cells at levels equivalent to wild type. This contrasts with our own findings that HaloTag knock-in cells demonstrate a 50% reduction in expression at the CXCR4 locus. We reproduce HaloTag knock-ins at the *TubA1B* locus in Hel 92.1.7 cells and find poor expression in these clones (approximately 4% compared to the significantly higher expression of mEos and mEGFP labelled *TubA1B* in these cells - all cells reported are heterozygous for insertion) (Figure S7,S8). Together these findings suggest a very gene specific effect of HaloTag knock-in at target loci.

Our interest is in the application of CRISPR mediated knock-in to generate high quality samples for robust quantitative SMLM. Clonal populations which are functionally validated and provide an established labelling density are potentially powerful quantitative tools. Therefore we perform single molecule imaging and cluster analysis of CRISPR and over-expression CXCR4 HaloTag cells to determine whether the changes in receptor clustering result-ing from ligand treatment can be established in these samples.

Our data shows that due to the sample to sample variation inherent to transfection, no significant change in cluster distribution is evident on ligand treatment. Conversely, our CRISPR cells consistently demonstrate significant differences in cluster statistics, consistent with the clustering and trafficking of CXCR4 on treatment with CXCL12. These results suggest that the generation of endogenously expressing samples are better suited to robust single molecule quantification.

## Conclusions

The use of CRISPR-Cas9 gene editing to introduce endogenously expressed fluorescent proteins remains a relatively small part of the literature, however with ever increasing knock-in efficiencies in classically difficult cell types, these experiments are becoming increasingly commonplace.

We demonstrate that targeting genes with identical guides, linkers, and homology templates but different tags achieves markedly different expression profiles and behaviours. To our knowledge, this has not been reported in the literature pertaining to CRISPR mediated knock-in to date. This variation in expression is likely to a myriad of regulatory factors which are, in our hands, gene specific. Our results show that mEGFP is consistently well expressed regardless of the gene targeted, however none of our mEGFP knock-ins are bi-allelic, suggesting that in our targets there is an upper limit to the amount of tagged, endogenous protein tolerated. This work is consistent with reports from the Allen Institute, which find that on generation of a large knock-in mEGFP library specific genes do not tolerate bi-allelic insertions(39). The abnormal stability of mEGFP tagged CXCR4 implies that this is likely due to a functional over-expression on mEGFP knock-in at the CXCR4 locus. In a recent review interrogating and discussing the effects of over-expression, Stepanenko and Heng highlight the genomic, epigenetic, and phenotypic effects of both transient and stable transfection on cells(14). CRISPR mediated labelling offers a number of distinctive advantages in advanced light microscopy, including SMLM. However, our results show that establishing the effects of knock-ins is critical to generating useful biological models. This is particularly important in light of both increasing applications of CRISPR for HDR, and the growing evidence of other important caveats to CRISPR, including off-target effects and low rates of correct HDR. This study suggests that there is a need to determine the properties of both fluorophores and knock-in sequences which may affect function and expression. While we explore monomericity and codon optimisation, a myriad of other factors including maturation rates and RNA stability are likely to be important. An ‘ideal’ insert is needed to fully realise the potential of these approaches, however CRISPR knock-in still promises to generate models which overcome classic limitations in cell biology and single molecule imaging. Finally, our results show that despite the complexity involved in the generation of CRISPR lines which express specialist reporters well enough for high quality SMLM, the result is a stable population suitable for high quality single molecule imaging. Compared to the equivalent transfected samples, CXCR4 knock-ins allow for accurate cluster quantification. As imaging approaches and analytical methods consistently improve in the single molecule field, the generation of robust samples through CRISPR is likely to be a valuable addition to the growing single molecule arsenal.

## Materials and Methods

### A. Cell culture and transfection

Human Embryonic Kidney 293T (HEK293T) cell lines were cultured in Dulbecco’s Modified Eagle’s Medium (DMEM) supplemented with Penicillin/Streptomycin (1%), 2mM L-Glutamine, and 10% Foetal Bovine Serum (FBS). Human Erythroleukaemia 92.1.7 cells (Hel 92.1.7) were maintained in identically supplemented Roswell Park Memorial Institute (RPMI) 1640 medium. All cells were cultured at 37°C and 5% CO_2_. Adherent cells were passaged by washing T75 cell culture flasks (Corning) twice with sterile Phosphate Buffered Saline (PBS) without calcium or magnesium, before incubation with Trypsin-EDTA (Thermo Scientific) for 5 minutes at 37°C. Cells were then re-suspended in complete DMEM before passaging at an appropriate dilution in a fresh flask. Suspension cells (Hel 92.1.7) were passaged by spinning cells at 1200 RPM for five minutes before re-suspension in fresh complete RPMI, dilution in fresh media, and incubation.

Adherent cells were transfected with Lipofectamine 3000 (Thermo Scientific) as per manufacturer guidelines. Briefly, cells were plated at a density of 1×10^5^ in 12 well plates (Thermo Scientific) 18 hours before transfection. Immediately preceding transfection, cells were washed twice with sterile PBS before incubation in Optimem. The px459 guide and Cas9-puro expressing plasmid was used (Addgene plasmid #62988 was a gift from Feng Zheng) to introduce the previously reported *TubA1B* targeting guide C or CXCR4 targeting guide with Cas9 elements to transfected cells(15, 40).

Suspension cells were transfected using the Neon electroporation system (Thermo). Briefly, cells were resuspended in a final total volume of 10*μg*l buffer R with 1*μ*g of total DNA (equimolar guide C px459 and donor) and a total of 1 × 10^5^ cells. 10 *μ*L tips were used with 2x pulses at 20ms pulse width and 1450V.

For over-expression studies, CXCR4-HaloTag CO sequence identical to that which is used in knock-in studies was cloned into a pcDNA3.1 over-expression vector and transfected at 25ng per 35mm MatTek dish. To maintain equivalent transfection efficiencies across experiments, transfection mixtures were bulked to a final total DNA concentration of 1*μ*g with empty pGem-T-Easy vector.

### B. Donor vector cloning

*TuBA1B* donor plasmids were generated by ordering gBlock synthetic DNA (Integrated DNA Technologies) for both left and right homology arms and each of the inserts tested (mEos 3.2, mEos 3.2 codon optimised, mEos 4b, mEos 4b codon optimised, HaloTag codon optimised and mEGFP). Codon optimisation was performed using IDT’s online tool. Each fragment was designed with a 20bp segment overlapping the adjacent arm. Homology arms and inserts were cloned into an empty pGem-T-Easy backbone using a HiFi Gibson Assembly kit (New England Biolabs).

2*μ*L of each 10*μ*L reaction volume was transformed into competent cells as described in the previous section. Individual colonies were selected after overnight growth on ampicillin plates, miniprepped (using manufacturer instructions included in the GeneJet kit Thermo Fisher), and subject to an EcoRI test digest to verify the presence of an insert. Clones carrying an insert of the correct size were further verified by Sanger sequencing. Validated plasmids were amplified.

Donor plasmids for CXCR4 genome engineering were generated by sub-cloning codon-optimised, mEos 4b, mEos 4b and HaloTag as well as mEGFP synthesised as doubled stranded DNA gBlocks (Integrated DNA Technologies) into the CXCR4 donor vector described previously using XhoI and XbaI restriction enzymes(40).

### C. Flow cytometry and single cell sorting

Validation and verification of knock-ins was performed using an Accuri C6 Flow Cytometer (BD Technologies). Samples were gated according to forward and side scatter, with positive fluorescence for these experiments detected in the FL1 channel. Single cell sorting was very generously performed by Matt MacKenzie (TechHub) on a BD FACSAria Fusion cell sorter with a 100*μ*m nozzle at 20 psi. Gates were set on the brightest population of cells to ensure that both highly expressing and correctly editing clones were selected.

### D. Quantitative RT-PCR

To examine the abundance of the endogenous mRNA expression levels in CXCR4 CRISPR-tagged and WT Hek293 cells, we carried out real time quantitative RT-PCR (qRT-PCR). Quantification of gene expression was performed using the ABI Prism 7500HT Sequence Detection System (Applied Biosystems, Foster City, CA, USA) and SYBRgreen master mix technology. Relative gene expression quantification was performed according to the comparative Ct method using GAPDH as an endogenous control. mEGFP-, Haloand mEos4b-tag specific fragments were amplified using a common forward primer to CXCR4 (5’-GAGTCTTCAAGTTTTCACTCC3’) and the following tag-specific reverse primers (mEGFP – 5’GCTTCATGTGGTCGGGGTAGC; Halo – 5’GAAGTACCCAAGGTCCGGTTTG-3’ mEos4b – 5’CGCGATTTCCATAGTGGAAGGC-3’. An endogenous CXCR4 fragment not specific to the tag was also amplified using Forward (5’-ATGGAGGGGATCAGTATATACAC-3’) and reverse 5’-GCCACTGACAGGTGCAGCCTGTAC-3’) primers.

### E. Western Blotting

Clonal populations were expanded in 6-well plates (Thermo Scientific) before lysis in NP-40 buffer with proteolysis inhibitor (Sigma). Lysates were prepared by 30 minutes incubation on ice in NP-40 with protease inhibitors (Sigma) before a final 10 minute spin at 14000 rcf. The supernatant was then added to 2x reducing sample buffer and boiled for 5 minutes.

Western Blots were prepared through SDS-PAGE on 4-12% gradient Bolt gels (Thermo Scientific). Gels were run for 15 minutes at 70 V, and 45 minutes at 125 V. Once run, each gel was transferred to a polyvinylidene difluoride (PVDF) membrane (Bio-Rad) using a Turbo transfer system (Bio-Rad). After transfer, membranes were blocked with 4% BSA in 0.1% Tween-20 Tris buffered saline (TBST) and probed with the relevant primary antibodies. Secondary incubations were performed using fluorescent antibodies (LiCor Instruments) in TBST. Anti-mouse 680 and anti-rabbit 800 fluorescent antibodies were used for detection using an Odyssey Fc (LiCor Instruments). Tubulin and GAPDH probes were used as previously described. Anti-CXCR4 antibodies Fusin C8352 (labelled antibody 1) and knock-out validated ab124824 (labelled antibody 2) were obtained from Thermo and Abcam respectively. Finally, Image Studio was used to quantify western blots.

### F. CXCR4 Functional Assays

Cells were transiently transfected according to the manufacturer’s instructions using FuGENE-HD transfection reagent (Promega, Wisconsin, USA) with plasmid cDNA coding for Gα i1/Nluc (synthesised by GeneArt, Invitrogen) and Venus/Gγ 2 (a kind gift from A/Prof Kevin Pfleger) 24 hours after seeding 350,000 cells/well in a 6-well plate. Cells were harvested with x 1 Trypsin-EDTA (Sigma Aldrich) and seeded into poly-D-lysine (Sigma Aldrich) coated white flat bottom 96 well plates (655089; Greiner Bio-One, Stonehouse, UK) at 30,000 cells/well 24 hours before performing an assay.

For each BRET assay, media was removed from 96well plates containing CRISPR/Cas9-modified or wildtype HEK293 cells and incubated with 1x HANK’s Buffered Salt Solution (1xHBSS; 25mM HEPES, 10mM glucose, 146mM NaCl, 5mM KCl, 1mM MgSO4, 2mM sodium pyruvate, 1.3mM CaCl2, 1.8g/L glucose; pH 7.45) supplemented with 0.1% BSA for 1 hour at 37°C under atmospheric CO_2_. Furimazine (Promega, Wisconsin, USA) was then added to a final concentration of 10 μM and incubated for 5 minutes before filtered light emissions where measured at 460 nm (80 nm bandpass) and 535 nm (60 nm bandpass) continuously at 37°C using a PHERAStar FS plate reader (BMG Labtech). After 5 reads CXCL12 (0.1 pM – 100 nM; Preprotech, Rocky Hill, USA) or buffer (HBSS containing 0.1% BSA) was added to triplicate wells and BRET was measured for a further 25 cycles. Raw BRET ratios were calculated by dividing the 535 nm emission by the 460 nm emission with the maximal change in BRET used for analysis.

For cAMP assays, wild type or CRISPR/Cas9-modified HEK293T were maintained as described for BRET assays above. On the day of assay cells were harvested with Dulbecco’s Phosphate Buffered Saline (DPBS) supplemented with 0.2g/L Ethylenediaminetetraacetic acid (EDTA; Sigma Aldrich) pre-warmed to 37°C and re-suspended in stimulation buffer (HBSS containing 500 *μ*M 3-Isobutyl-1Methylxanthine and 0.1% BSA) at 400,000 cells/ml. cAMP production was measured using a LANCE^®^ cAMP Detection Kit (PerkinElmer) following the manufacturer’s instructions. Briefly, 5 *μ*L of cells were added in duplicate to a white 384 well plate (ProxiPlate-384, PerkinElmer) containing buffer only (stimulation buffer containing Alexa Fluor^®^ 647-anti cAMP antibody) or Forskolin (0.5 *μ*M, Sigma-Aldrich) in the absence or presence of CXCL12 (1 pM 1 *μ*M) for 45 minutes at room temperature. The reaction was then stopped by addition the cAMP detection buffer containing Eu-W8044 labeled streptavidin and biotin-cAMP. Plates were then incubated for 1 hour at room temperature and fluorescence was measured at 615 nm and 665 nm, respectively, 50 *μ*s after excitation at 320 nm using an EnVision 2102 microplate reader (PerkinElmer).

### G. Imaging (PALM and dSTORM)

Clones of interest were imaged on 35mm MatTek Dishes (MatTek Corporation) on a Nikon N-STORM system (Andor iXon Ultra DU897U EMCCD, Ti-E stand, Perfect Focus, Agilent MLC400 laser bed). A 100x 1.49 NA TIRF Objective was used for each acquisition, which was then reconstructed using ThunderSTORM (41) (Maximum Likelihood, Integrated Gaussian PSF fitting).

Hel 92.1.7 cells were treated with phorbol 12-myristate 13acetate (PMA) (Sigma) and thrombopoeitin (TPO a generous gift from Ian Hitchcock) overnight to drive spreading and differentiation prior to imaging. Once seeded on MatTek dishes, cells were washed twice with PBS before treatment with microtubule stabilising buffer (MTSB 80mM PIPES pH 6.8, 1mM MgCL_2_, 4mM EGTA) and 0.5% Triton-X100 for 30 seconds before fixation with ice cold Methanol at −20°C for 3 minutes. Samples were then washed with TBST and imaged in PBS for CRISPR-PALM.

CXCR4 knock-ins were labelled with Janelia Fluor 549 (generously provided by Lavis lab) prior to fixation(42). Cells were plated on 35mm MatTek dishes overnight before the addition of 250nM JF549 in complete DMEM and a subsequent 15 minute incubation. Samples were then washed with PBS and complete media before a further 30 minute incubation in unlabelled complete media. Finally, cells were washed 3x with PBS before fixation in formalin for 10 minutes. For CXCL12 treated samples 100ng (10*μ*M) recombinant human CXCL12 (SDF1*α*, Thermo) was added for 4 minutes prior to fixation. Finally, these samples were imaged in blinking buffer (100mM mercaptoehylamine-HCL, 1g/mL catalase, 50g/mL glucose oxidase, PBS) at 20ms exposures and approximately 150kW laser intensity (with low level 405 illumination added on successfully shelving JF549 into a dark H Cluster analysis state).

### H. Cluster analysis

Images were reconstructed as previously described by Khan *et al.* using ThunderSTORM(41, 43). Prior to cluster analysis, images were drift corrected (cross correlations), and filtered to remove detections with uncertainties above 75nm. Duplicates were removed, and blinks recurring within 50nm in successive frames were merged. Filtered detections were clustered using persistence based clustering, implemented using RSMLM and KNIME (workflow available on request) (44, 45). Detection density was estimated by counting the number of neighbouring detections within 20nm and the persistence threshold was set to 10 detections. For each field of view the central 10 *μ*m^2^ was cropped and analysed. Clusters with less than 10 detections were removed from the analysis. Cluster area was defined by the convex hull of the detections within a cluster.

### I. Image and statistical Analysis

Statistical analysis was performed using GraphPad PRISM 6, with statistical tests as indicated in figure legends. Briefly, significance was determined using either 2-way ANOVA or 1-way ANOVA with multiple comparisons. *n* = 3 across samples unless otherwise stated, and error bars represent standard deviation of the mean. FRC measurements were performed using the recently published NanoJ-SQUIRREL plug-in(46, 47).

CXCR4 functional data were normalized to the maximal response generated by wildtype HEK293T cells and analysed using GraphPad Prism 7. Concentration-response data were fitted with sigmoidal curves generated using non-linear regression assuming a slope of 1. Statistical analysis was performed using GraphPad Prism 7 using a one-way ANOVA and Dunnett’s multiple comparisons test. Number of individual repeats performed are indicated in the figure legends. P<0.05 was considered significant.

## AUTHOR CONTRIBUTIONS

AOK devised project and experiments, performed transfections, single cell sorting, cloning, western blotting and single molecule imaging. CW designed and generated CXCR4 constructs and functional assays for knock-in Hek293T, as well as input to manuscript and experimental design. JAP performed image analysis of single molecule data and revised manuscript. JY and AS assisted with cloning of constructs. SH, NSP, SGT, and NVM supervised project and revised manuscript for submission.

## Acknowledgements

The work in the author’s laboratories was supported by the British Heart Foundation (PG/13/36/30275; FS/15/18/31317, PG/16/103/32650). This work was in part supported by a Team Science grant from COMPARE as well as fellowship support to CWW from the National Health and Medical Research Council (NHMRC) of Australia (CJ Martin Research Fellowship, 1088334) and from a University of Western Australia Fellowship Support Grant to CWW and an MRC program grant to SJH [MR/N020081/1]. We thank A/Prof Kevin Pfleger for the Gα i1/Nluc and Venus/Gγ 2 constructs and Lavis Lab for the generous provision of Janelia Fluor for this project. The authors would like to acknowledge Professor Steve Watson for his ongoing support, and Drs. Kabir Khan and Jasmeet Reyat for their insightful comments on the manuscript. The authors would also like to acknowledge the Imaging Suite at the University of Birmingham for support of imaging experiments. Imaging facilities used in this project were funded by COMPARE. Finally the authors would like to thank the TechHub cell sorting facility, in particular Matthew MacKenzie.

## Supplementary Note 1: Supplementary Figures and Tables

**Fig. S1.**
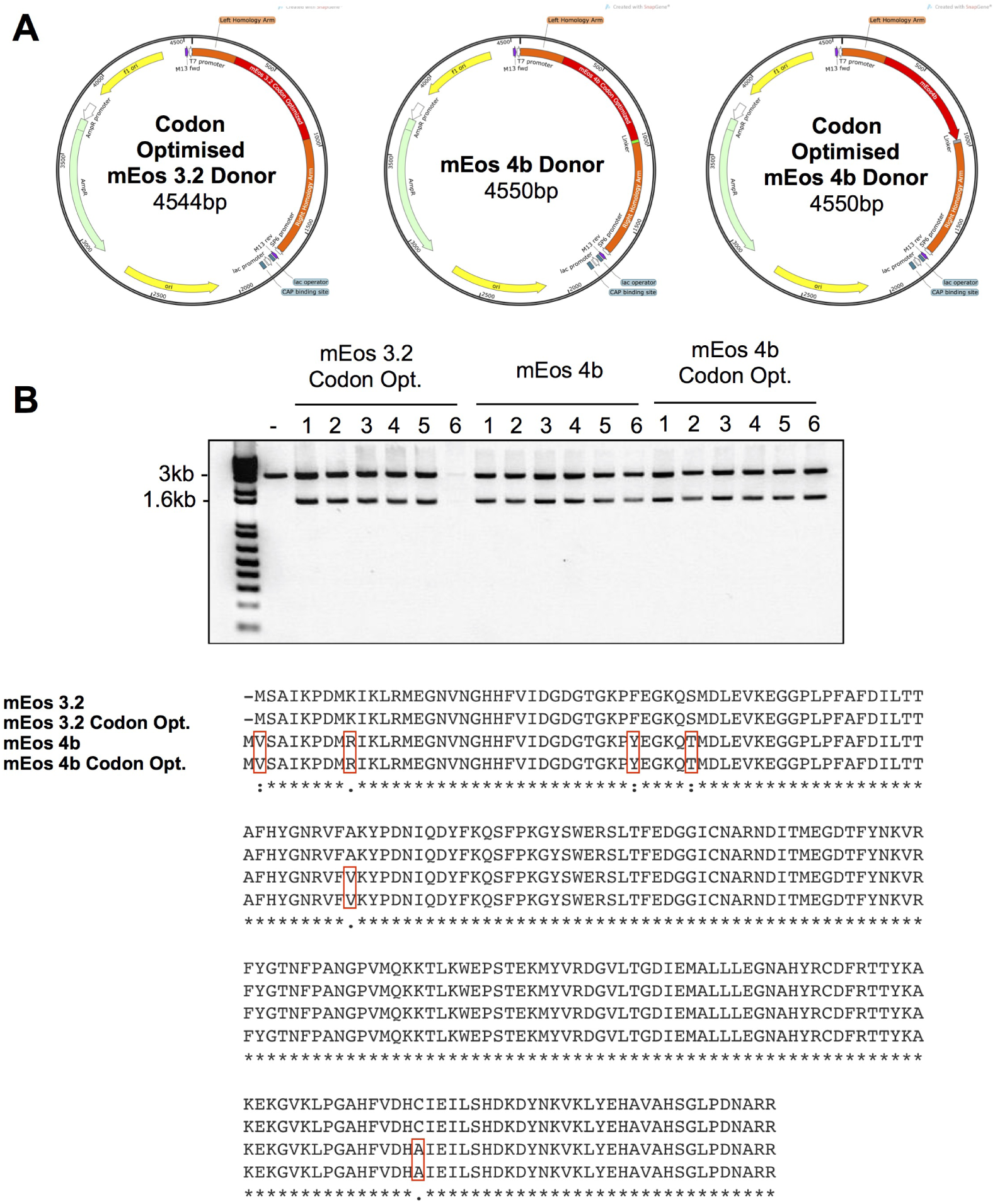
Generation of mEos variant *TubA1B* donors. (A) Donor plasmids with identical linkers and homology arms harbouring mEos variants were designed. These include a codon optimised mEos 3.2 sequence, as well as a truly monomeric version of the protein reported as mEos 4b, and a respective codon optimised variant. (B) gBlocks were cloned into pGem-T-Easy backbones and validated through EcoRI digest. Negative controls are 3000bp in length, while successfully cloned vectors demonstrate an insertion which runs at approximagely 1600bp. Finally sequence alignments show differences in the 3.2 and 4b protein sequence, and that codon optimisation does not alter the reading frame or coding sequence reported. Amino acids marked in red are where the mEos 4b protein differs from its predecessor mEos 3.2.

**Fig. S2.**
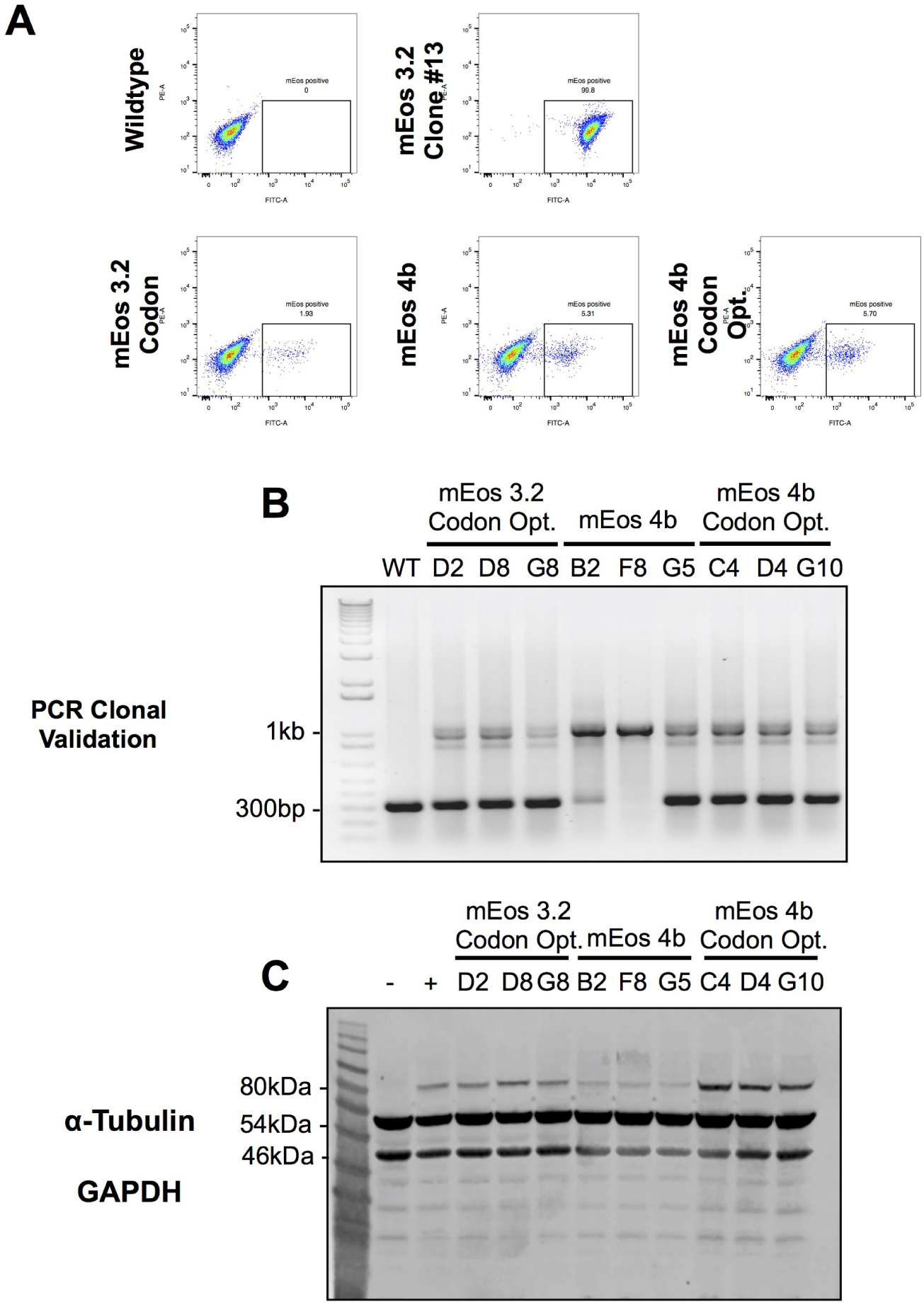
Generation, sorting, and genomic validation of Hel 92.1.7 *TubA1B* CRISPR clones. (A) Hel 92.1.7 cells co-transfected with a guide targeting *TubA1B* and donor templates carrying codon optimised mEos 3.2, mEos 4b, and codon optimised mEos 4b were single cell sorted into a 96 well plate. (B) Clones were expanded and validated by PCR. The amplification of wild type DNA results in a 300bp band, while heterozygous clones positive for fluorophore insertion at the *TubA1B* locus also feature a second band at approximately 1000bp.(C) Full unedited western blot of gels presented in figure 2

**Fig. S3.**
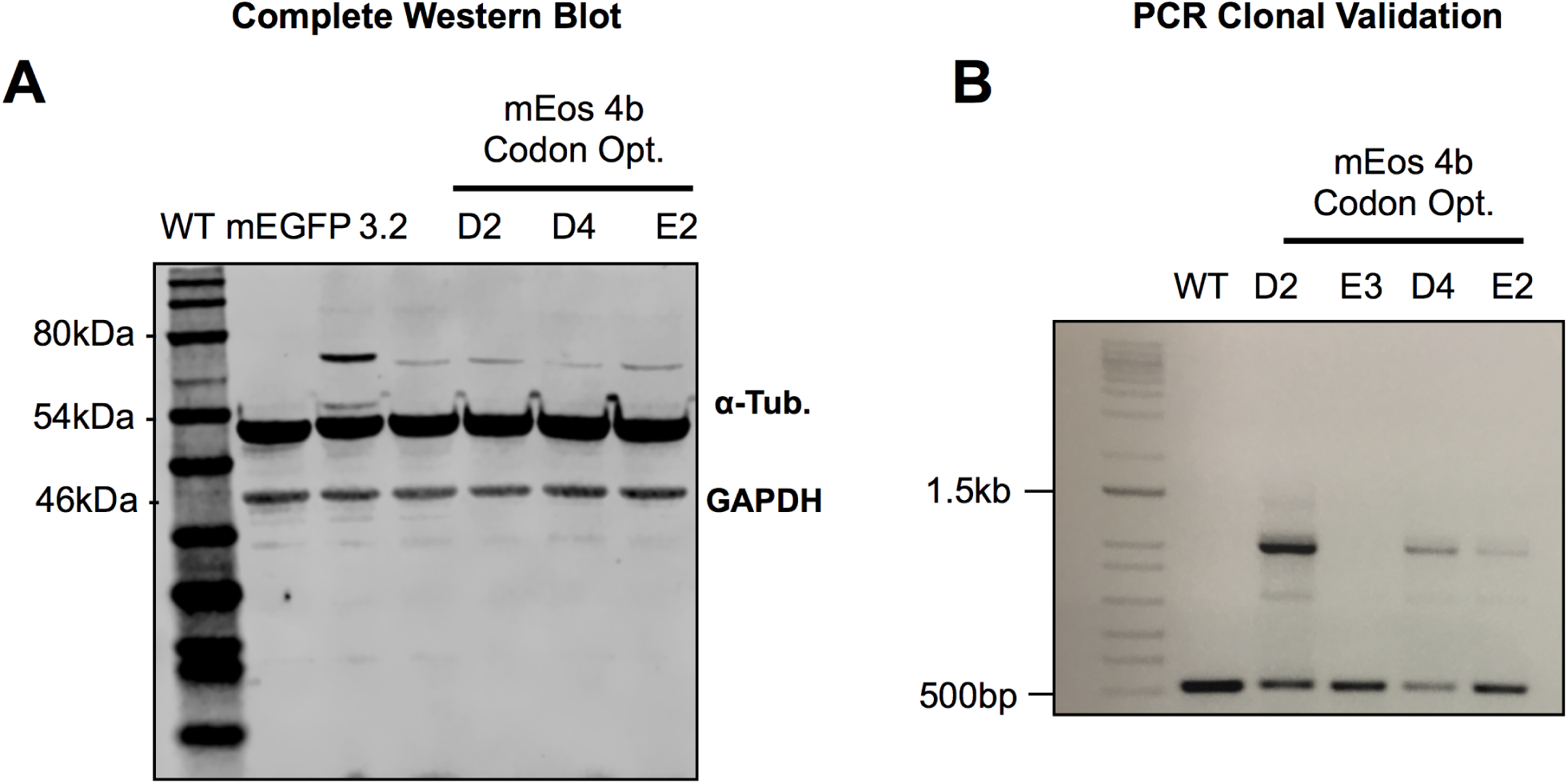
Complete gels from main text. (A) Full gel of Hek293T *TubA1B* clones reported in figure 2. (B) PCR validation of mEos 4bCO clones, compared to wild type, clones D2, D4, and E2 demonstrate a band approximately 700 base pairs heavier than the wild type sequence, indicating a successful heterozygous insertion of the sequence at this locus.

**Fig. S4.**
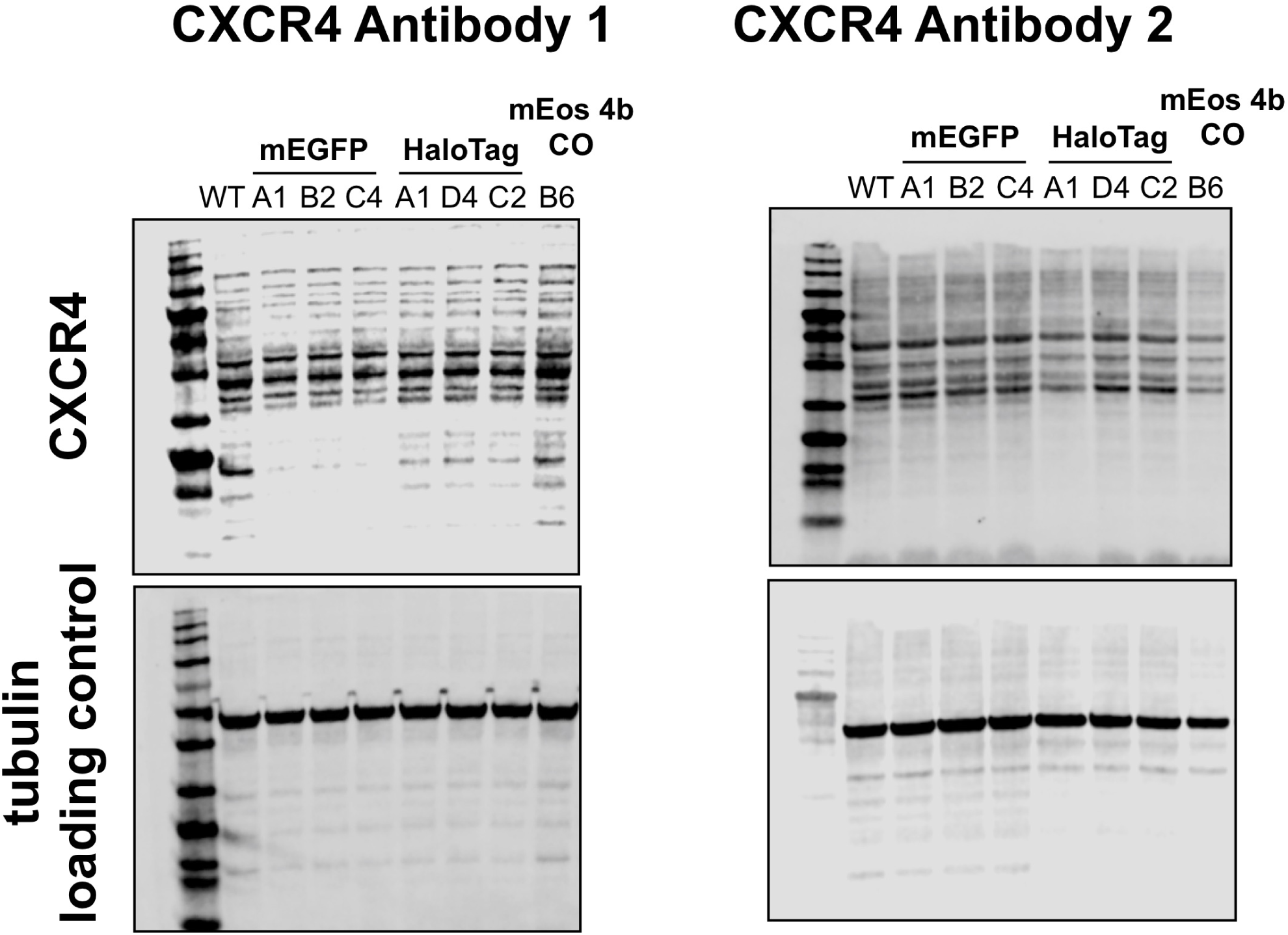
Western blots of CXCR4 knock-in and wild type Hek293T. Poor detection of CXCR4 with both a knock-out validated and a fusin antibody was observed despite detection and validation by imaging, RT-PCR, and functional assays.

**Fig. S5.**
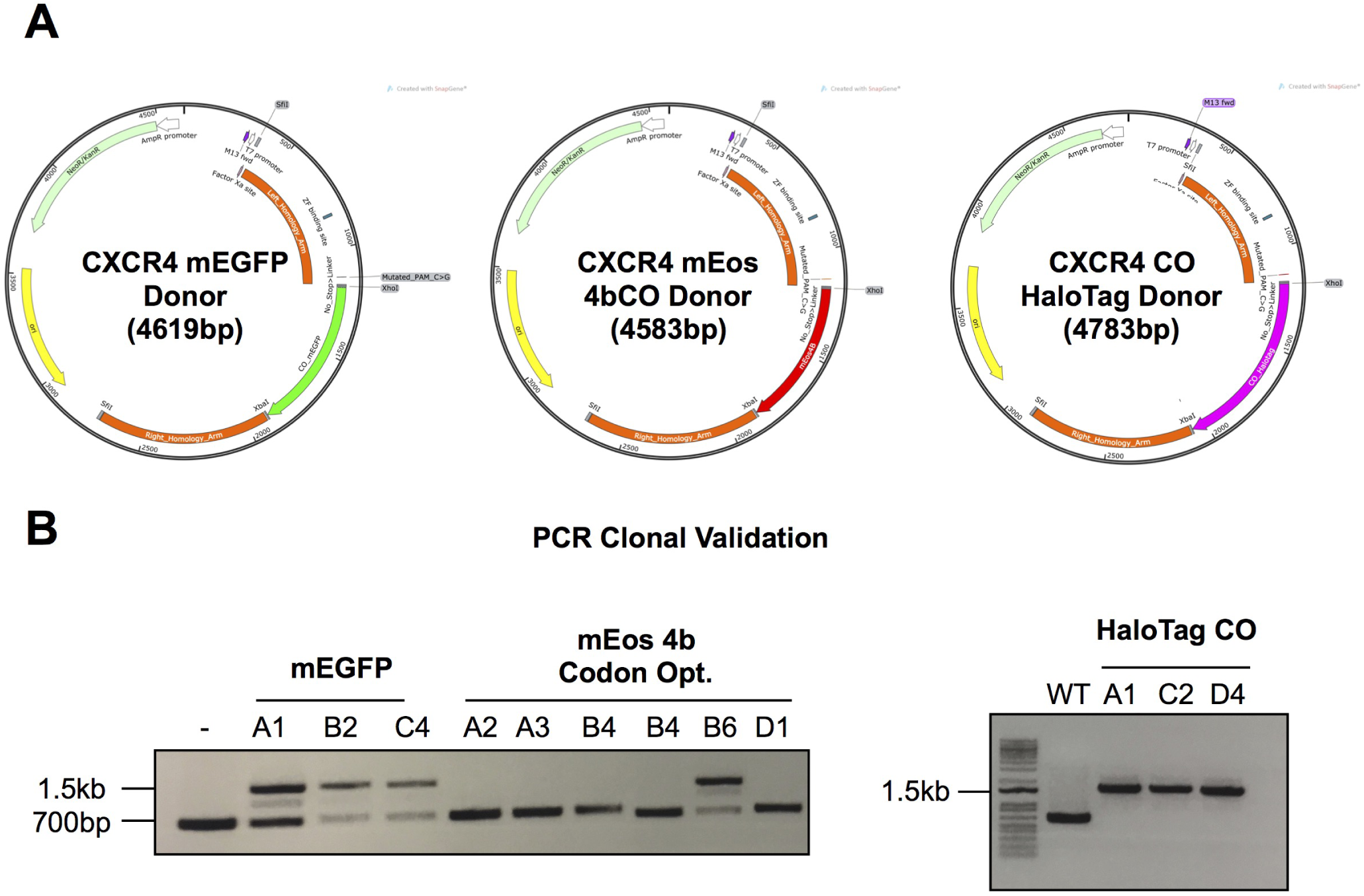
Design of CXCR4 donor plasmids and genomic validation on knock-in and clonal selection. (A) HDR donor plasmids targeting the CXCR4 C-terminus were generated following the design and cloning strategies previously described by White *et al.*. Donor plasmids carrying mEGFP, mEos 4b CO, and a codon optimised HaloTag sequence were designed and validated by sequencing. (B) On transfection and single cell sorting, cells were validated by PCR to determine the correct insertion of donor sequence at the target loci. mEGFP clones were successfully validated as heterozygous, while mEos 4b CO cells were generally negative due to the low level fluorescence observed (as shown in figure 3, mEos 4b CO appears to agglomerate in the validated clones). HaloTag CO knock-ins proved homozygous for insertion, with a loss of the wild type sequence in the three clones tested.

**Fig. S6.**
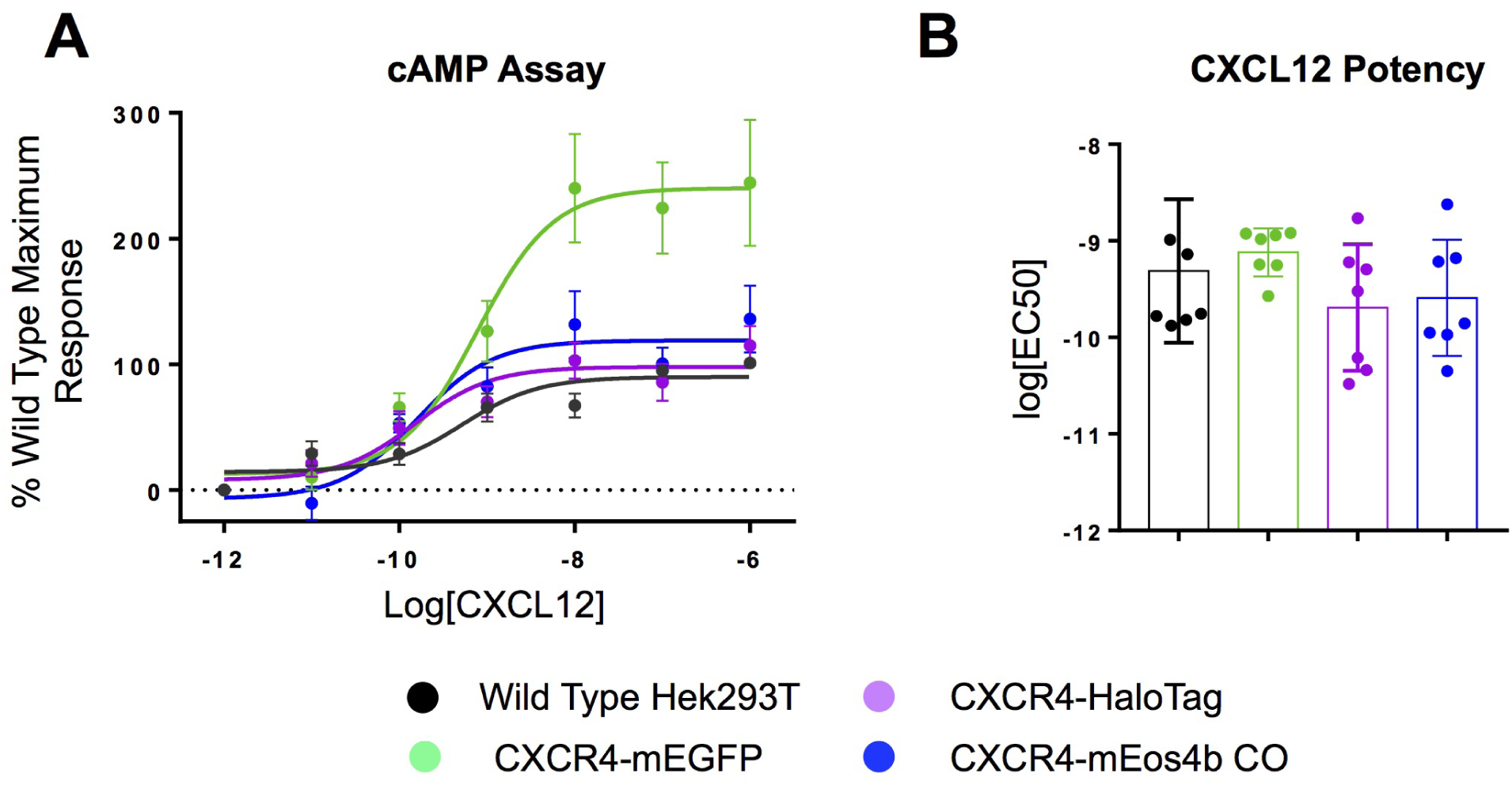
Functional validation of CXCR4 knock-in clones through cAMP assay. (A) Modulation of cAMP accumulation mediated by CXCL12 (1 pM – 1 *μ*M) in HEK293 cells expressing wildtype or genome-edited CXCR4. (B) Potency of CXCL12-mediated g protein activation in cells expressing wildtype or gene-edited CXCR4. Points and bars represent % maximum CXCL12 response observed in wildtype HEK293 cells ± S.E.M. of seven independent experiments.

**Fig. S7.**
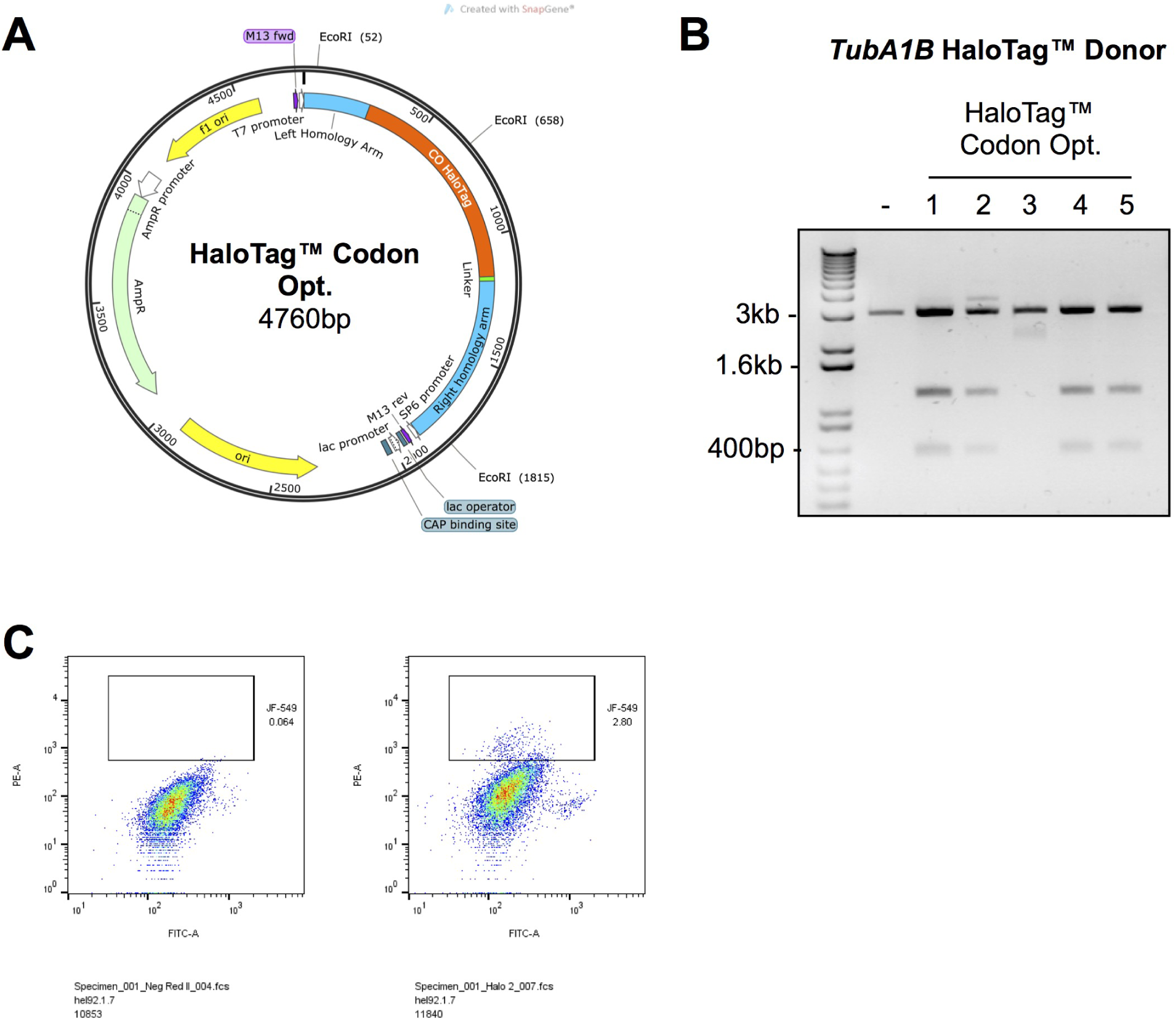
Generation of *TubA1B* HaloTag CRISPR donor vectors. (A) The *TubA1B* donor template was sub-cloned to include a donor HaloTag sequence as previously described. (B) Vectors were prepped and validated by EcoRI digest before sequencing, with successfully cloned vectors demonstrating two additional bands compared to the negative linearised protein. (C) Hel 92.1.7 cells were co-transfected with the validated HaloTag *TubA1B* donor and a targeting guide, single cell sorting was thenperformed after incubation with Janelia Fluor 549, selecting for the brightest cells as in previous experiments.

**Fig. S8.**
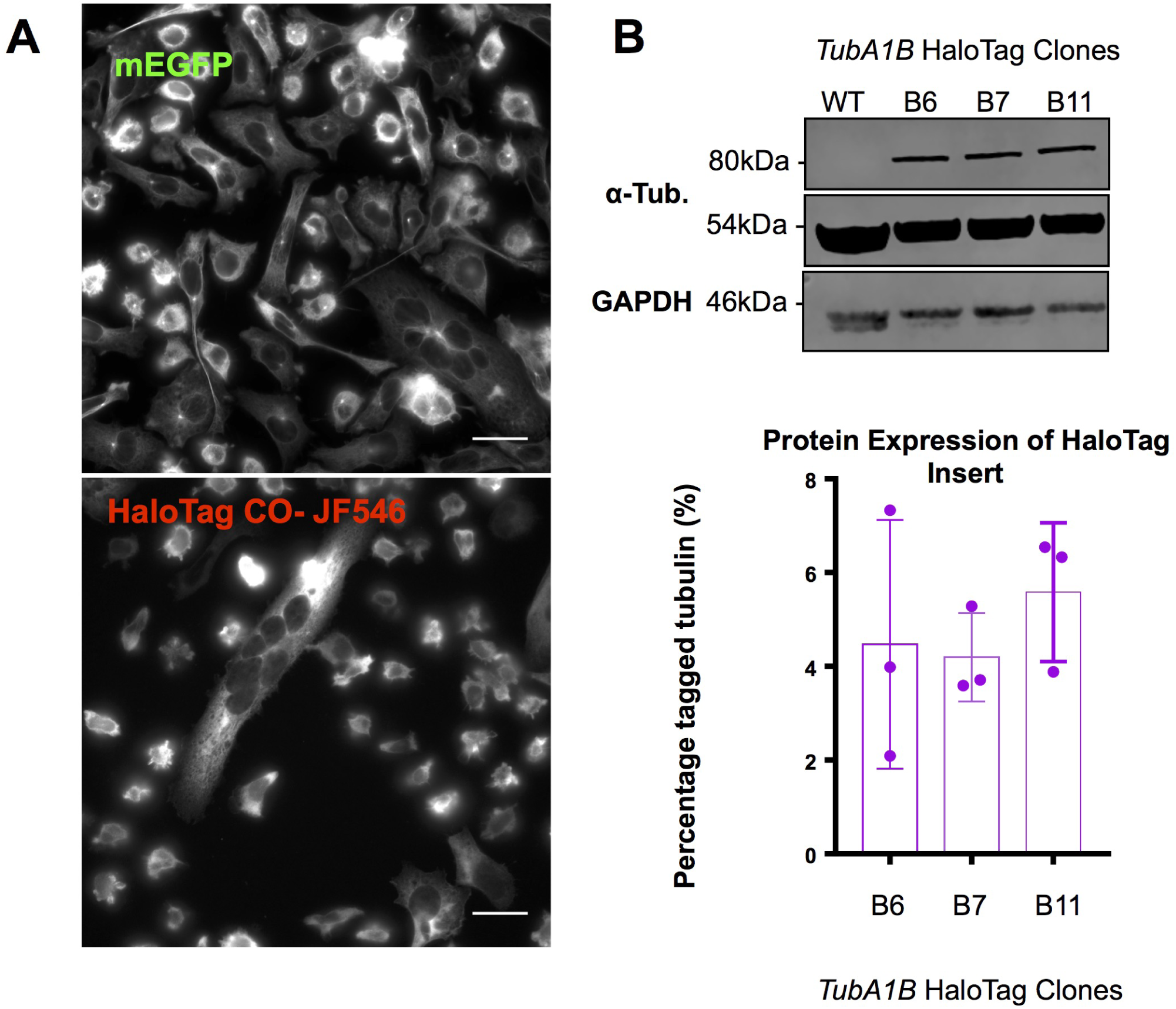
Validation of HaloTag-*TubA1B* knock-in clones. (A) Hel 92.1.7 cells positive for the HaloTag insert were compared to mEGFP labelled *TubA1B* in the same cell line, while HaloTag labelled cells are fluorescent on incubation with JF549 and show microtubule distribution, brightness in these cells is substantially lower than their mEGFP counterparts. (B) Quantification of the expression level of HaloTag *TubA1B* shows a substantially reduecd expression of the tagged allele compared to previously reported knock-ins at approximately 5% compared to 20% mEGFP and 18% mEos 4b CO as previously reported.

